# Stress-free state in human carotid arteries cannot be revealed without layer separation

**DOI:** 10.1101/2024.10.15.618414

**Authors:** Anna Hrubanová, Ondřej Lisický, Ondřej Sochor, Zdeněk Bednařík, Marek Joukal, Jiří Burša

## Abstract

Residual stresses are considered as a significant factor influencing the stress-states in arteries. These stresses are typically observed through opening angle of a radially cut artery segment, often regarded as a primary descriptor of their stress-free state. However, the experimental evidence regarding the stress-free states of different artery layers is scarce. In this study, two experimental protocols, each employing different layer-separating sequences, were performed on 17 human common carotid arteries; the differences between both protocols were found statistically insignificant. While the media exhibited opening behaviour (reduced curvature), a contrasting trend was observed for the adventitia curvature, indicating its closing behaviour. In addition to the different bending effect, length changes of both layers after separation were observed, namely shortening of the adventitia and elongation of the media. The results point out that not all the residual stresses are released after a radial cut but a significant portion of them is released only after the layer separation. Considering the different mechanical properties of layers, this may significantly change the stress distribution in arterial wall and should be considered in its biomechanical models.

## 1. Introduction

Biomechanical investigation of the presence of residual stresses (RSs) in arteries dates back several decades (1–3). Arterial vessels, whether in situ or post-extraction, cannot be deemed stress-free due to the existence of RSs in the vessel walls (4,5); they are manifested by the typical opening of a circumferential strip of the artery wall documented in various experimental studies (4,6). Although absolute values of RSs may be smaller in comparison to the stresses experienced by arterial wall *in vivo*, their significance in shaping the resulting stress distribution should not be overlooked. The strain stiffening response of arterial tissue amplifies the impact of RSs, rendering the stress distribution more uniform throughout the wall thickness (5–8).

Patient-specific computational models of pathological arteries have confirmed their capability to predict arterial tissue rupture. For instance, Polzer et al. (9) demonstrated better rupture risk prediction of abdominal aortic aneurysm (AAA) using a novel probabilistic rupture risk index, outperforming the current treatment guidelines (10). A similar predictive approach could be applied to assess the vulnerability of atheromas, particularly those located in life-threatening locations such as coronary or carotid arteries. The inclusion of RSs in computational modelling is necessary, as the initial stress-free state is a prerequisite for any computational model (11). However, multiple studies seem to overlook the effect of RSs and strains in atherosclerotic carotid arteries (12–17) when evaluating their mechanical response to *in-vivo* mechanical loading. Although efforts have been made to incorporate the RSs into finite element (FE) models (11,18–22), proving the importance of RS in the resulting stress distribution, the scarcity of experimental data demonstrating RSs in the simulated artery types remains a challenge.

Most experiments focus on the entire artery wall without specific layer separation, thereby neglecting the multi-layered heterogeneous structure of the arterial wall. An experimental and computational study involving the complete human and porcine carotid artery bifurcation (5) affirmed that the inclusion of RSs in the computational models leads to a more uniform stress distribution and reduces maximal stress values on the inner wall surface. Greenwald et al. (4) explored different opening angles in two layers of bovine carotid arteries, noting larger opening angles in the inner layers compared to the outer ones. Holzapfel et al. (6) experimentally examined layer-specific residual deformations in human aortas, specifically exploring the opening angles of axial and circumferential strips from intima, media and adventitia layers. The methodology was further refined in (23) and applied to human carotid arteries in (24), where the adventitia (A) and media + intima (MI) layers were investigated separately. They recorded significant curvature changes after having released residual stretches in both axial and circumferential directions for both A and MI layers.

However, a deeper experimental investigation of this phenomenon is essential to gather data applicable as input for simulations of stresses in arteries with atheroma, as done, for instance, by Pierce et al. (19) and Sigaeva et al. (20). Approaches based on stress homogenization throughout the wall thickness, as seen in (25), may falter in highly non-homogeneous arteries with atheromas. The review (26) focusing on atherosclerosis does mention RS yet exclusively for the non-atherosclerotic tissue.

Hence, the objective of this study is to compare different experimental procedures ensuring credible experimental data on RSs demonstrated by residual deformation in separate layers, specifically in the circumferential direction, for healthy carotid arteries. Such insights can prove invaluable for computational modelling of arteries afflicted with atheroma and predictions of atheroma vulnerability.

## 2. Materials and Methods

For the purpose of this study, 17 common carotid arteries (CCAs) were harvested (between 1.4.2021 and 31.3.2024) during autopsies at Masaryk University, Department of Anatomy, with the approval of the local Ethics committee. Informed written consent from the body donors was obtained in Anatomy Bequest Program years prior to their death. The donor cohort comprised 8 males and 7 females, with an average age of 80.7±7.5 years. The samples were preserved frozen in a saline solution (0.9% NaCl) at −20°C since it was not possible to test the samples immediately after extraction. Some studies suggest that freezing may change the mechanical properties (27–30), however the change was observed in longitudinal direction only or at higher strains (28,29), or was mostly statistically insignificant (30). Other studies lean towards the conclusion that freezing at −20°C does not alter mechanical properties (31–34) and is actually better compared to mere refrigeration (30–32). On the testing day, they underwent a gradual thawing process and were stabilized in the saline solution, maintaining the temperature of 36.7±0.5°C. The samples did not outreach type IV lesion, as per the lesion characterization according to (35), showing none or mild signs of atherosclerotic changes.

### 2.1. Sample preparation and experimental protocol

The loose connective tissue was meticulously removed from the outer adventitia surface. Due to the limited sample length, varying between 10 and 30 mm, we were unable to obtain both circumferential and axial strips; therefore, only circumferential rings were extracted. Two distinct experimental protocols were implemented (see Fig. 1).

**Fig. 1:**
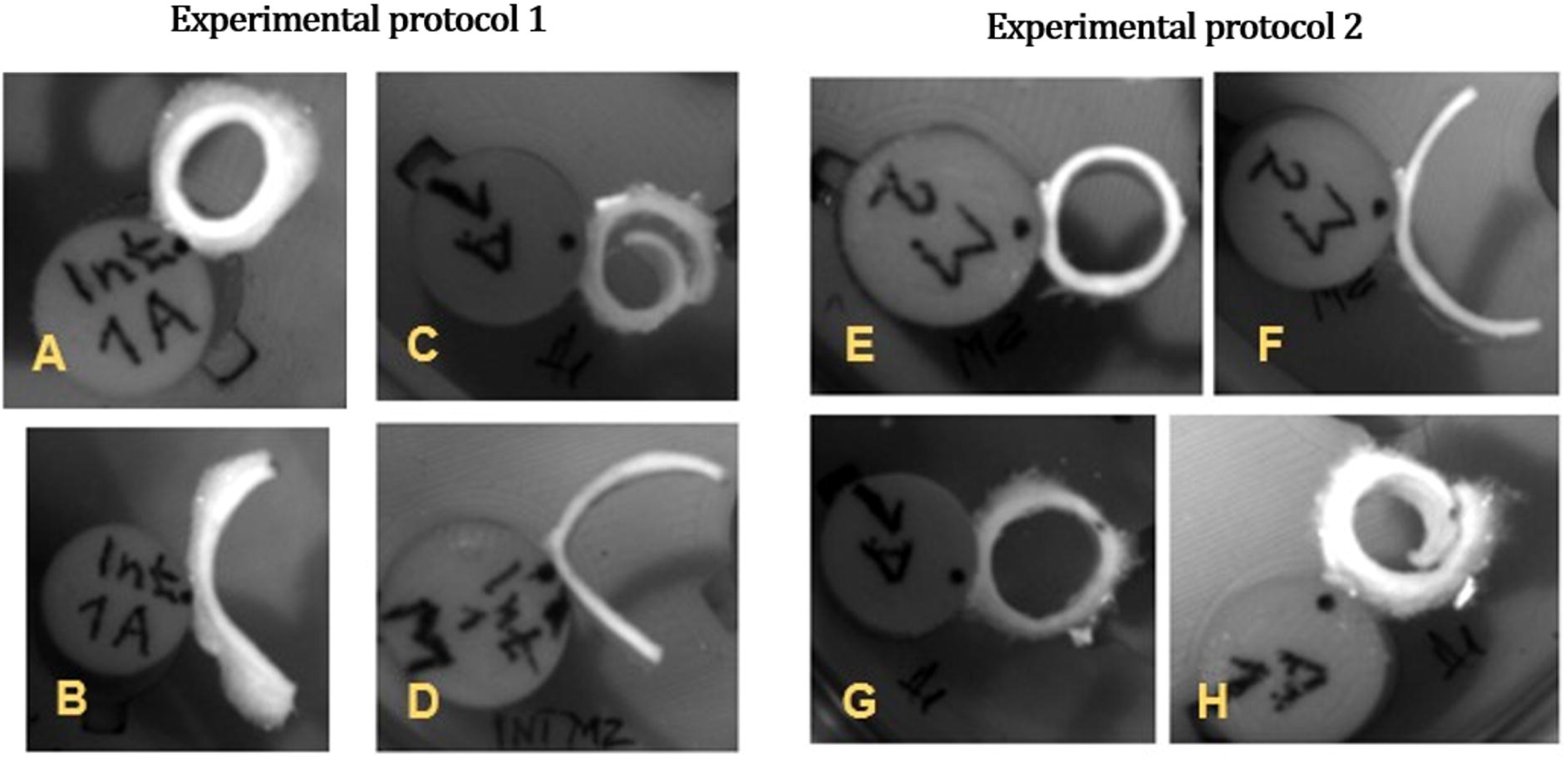
Two distinct experimental protocols were employed in this study to investigate the residual deformations of carotid wall layers and to compare different RS releasing procedures. The A (blue) and MI (yellow) layers were recorded within 22 hours after being radially cut. Black circles represent plastic cylinders used for specimen fixation.

The experimental protocol 1 (replicated from (6)) started with cutting the “intact” artery samples into 2 mm height circumferential rings, followed by equilibrating them in a 37°C saline solution for 30 minutes. Subsequently, they were glued pointwise to a plastic cylinder using cyanoacrylate adhesive, followed by a radial incision. This incision induced gradual opening of the rings, manifesting thus a release of circumferential RS, tensile at the outer and compressive at the inner artery surface. After 16 hours, the opened configuration was documented, and the adventitia was separated from the MI layer. Another recording occurred after additional 6 hours in the tempered bath.

In the experimental protocol 2, the A and MI layers were separated first. Attempts to isolate the intima from the media without damage were hindered by the natural thinness of the carotid intima. Moreover, the initial atherosclerotic stages related to patients’ age also blur the intima-media interface. Similarly to other studies (24), the inner layer was denoted as the MI layer. Then 2 mm circumferential strips were cut from both A and MI layers and glued pointwise to a cylindrical plastic tube. Images [<u>dataset</u>] were captured using a CCD camera (resolution 1280×1024) perpendicular to the tested ring 30 minutes, 16 hours and 22 hours after the radial incision, aligning with the intervals applied in (6) and maintaining time consistency across both protocols.

The thickness of the unseparated ring and of both layers was measured by a dial indicator at three locations for each specimen, the average value was considered for the subsequent analysis. The layer separation was examined by histological analysis (Fig. 2C, D) (standard haematoxylin and eosin stain) and quantified by the thickness ratio of the separated MI and A layers. A total of 22 intact specimens were prepared for protocol 1, yielding 21 adventitia and 20 media specimens; for the second protocol, 27 media samples and 25 adventitia samples were prepared. The specimen discard was due to occasional tissue damage and unsuccessful layer separation. The adventitia layer, distinguishable from the media by colour and texture, can be easily separated in a “turtleneck fashion” (24) (Fig. 2A). The media layer is stiffer and maintains its cylindrical shape even post-separation, whereas the adventitia becomes flat under gravity (Fig. 2B). It was observed that upon repeated immersion in the saline solution, the adventitia layer regains its cylindrical shape, demonstrating its high compliance.

**Fig. 2:**
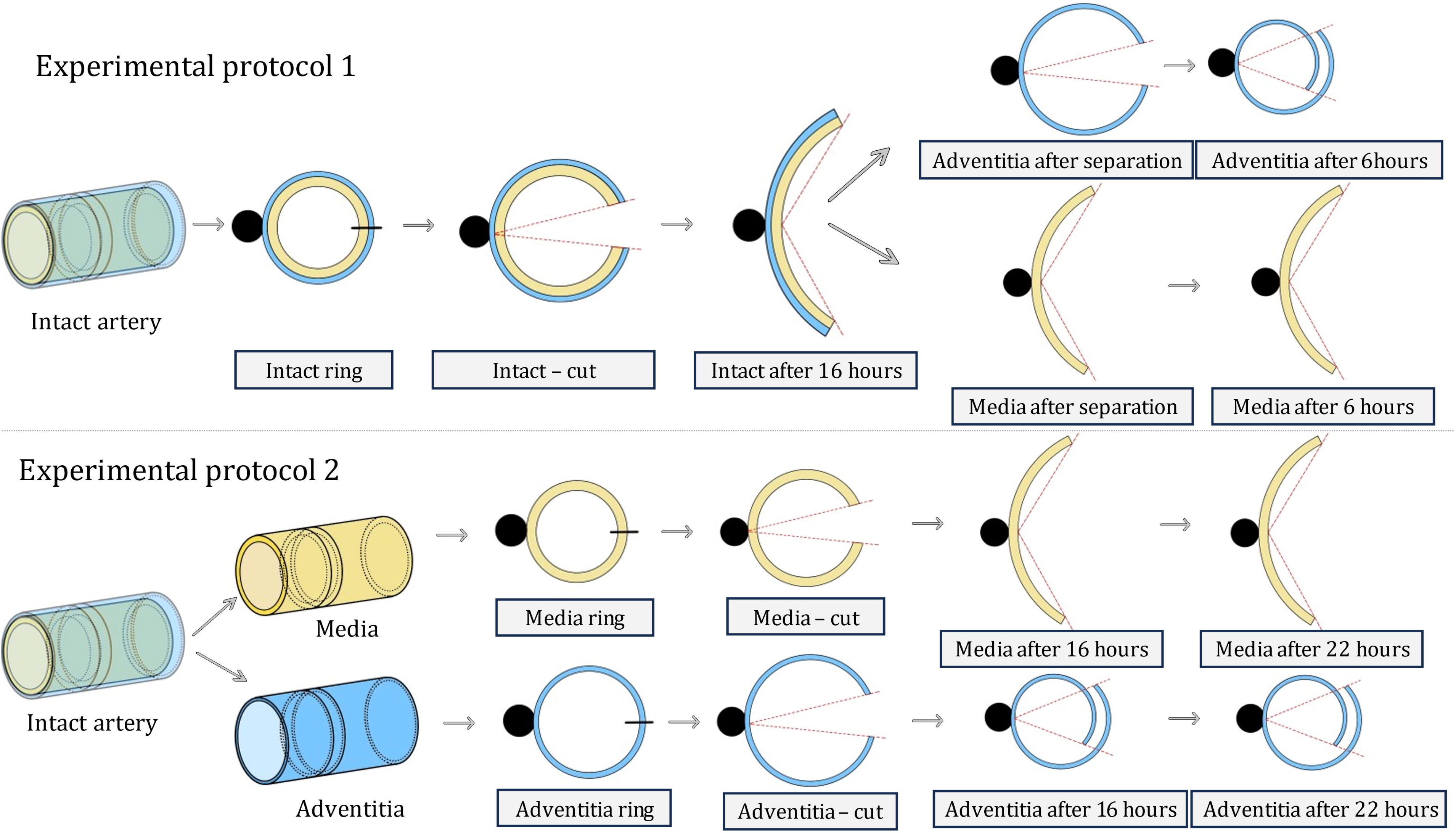
The adventitia and media separation in a “turtleneck fashion” (A). The adventitia layer turned inside-out for removal of the remaining media (B). Histological examination of adventitia (C) and whole artery wall (D) showed some remaining portions of the media on the adventitia.

Given the temperature-dependent nature of arterial mechanical responses (36,37), specimens were kept in the 37°C saline solution throughout testing for both experimental protocols, preventing tissue dehydration.

### 2.2. Image processing

Each acquired specimen image underwent analysis using an in house semi-automatic program in Python to obtain essential geometric parameters. The opening angle *Φ* [°] was defined by two lines pointed from the middle of the inner boundary to both ending points (equidistant). The definitions of the opening angle vary among studies, but they can be recalculated (8) for comparison, assuming the circular shape. However, local curvature represents a better RS quantifier independent of the strip length and asymmetry of the specimens (24); this is important especially for non-homogeneous atherosclerotic CCA walls. For this purpose, the inner and outer artery borders were manually traced, ensuring enough points (approx. 20 for each) for their accurate description. Subsequently, B-splines were utilized to approximate these traced borders and divided by 40 equidistant points. The sum of lengths between these points was considered as the approximate specimen length. Image scale was obtained using the known diameter of the plastic cylindric tube. Local curvature [mm^−1^] was calculated at each of the points using equation (1):

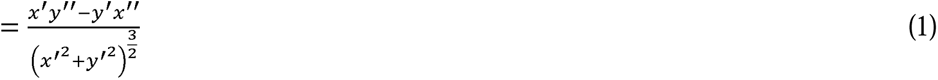

where *x*′, *y*′ and *x*″, *y*″ are the first and second derivatives of the B-spline, respectively. Subsequently, a local radius of curvature [mm] was calculated as follows:

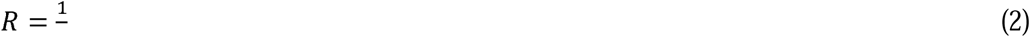

which was then averaged across all the calculated points within each segment boundary. Therefore, we evaluated local curvatures of the B-spline of the respective line (38 points for each B-spline); the averaged value represents then a global characteristic.

In summary, the six parameters of interest for both detached layers included their thickness, inner and outer curvature, inner and outer length, and opening angle.

### 2.3. Statistical analysis

Statistical analyses were conducted using the Minitab software. The Anderson-Darling normality test was employed for each dataset to ascertain a normal distribution of the data. If the normal distribution was confirmed, the data was presented as mean ± standard deviation; otherwise, the median and interquartile range were used. To assess differences in thickness, curvature and the opening angle of layers between both experimental protocols, a 2-sample t-test was applied when normal distribution was confirmed; otherwise, the Mann-Whitney test was employed. For variables measured within the same sample (e.g. length before and after the release of RS), the paired t-test was used for normally distributed data, and the Wilcoxon signed-rank test served as the nonparametric alternative.

Furthermore, Spearman’s correlation coefficient *ρ* was used for identification of a potential relationship between the opening angle and sample thickness. In all cases, statistical significance was considered if p < 0.05.

## 3. Results

### Thickness

Histological examination indicated that achieving a perfect separation of the adventitia layer was challenging as some portions of the media consistently adhered to the adventitia layer (Fig. 2C). Despite meticulous efforts, achieving a flawless separation was deemed unattainable even under the supervision of an experienced surgeon from St. Anne’s University Hospital. As the media consists of multiple very thin elastin membranes, they enable different separation planes and are prone to tearing (Fig. 2D). Despite the efforts to clean the media from the adventitia post-separation, the thickness variations of the adventitia layer persisted across specimens pointing out the inability of cleaning the retained media completely. A 2-sample t-test indicated no significant difference in thickness of the specific layers (MI and A) between both protocols 1 and 2. The mean thickness values for MI, A and the intact artery wall (I) were 0.60±0.14 mm, 0.36±0.11 mm and 0.92±0.15 mm, respectively.

### Opening angle

The opening angles for both experimental protocols are shown in Fig. 3, displaying values for detached layers as well as intact artery walls (for protocol 1). Predominantly positive values were observed for the media and even slightly higher for the intact wall. Conversely, negative values predominated for the adventitia, indicating strip closure rather than opening (Fig. 7C or H). Absolute values of both opening and closing angles increased in time for both layers and experimental protocols, which were compared for the separate layers in the final states. For both layers, the statistical analysis revealed no significant difference between the two protocols, therefore they were merged; the median of MI opening angle was 42.17° (13.38, 97.27), and the mean opening angle of A was −5.19±30.11°. For the intact wall, i.e. protocol 1 after 16 hours, the opening angle was 53.05±35.84°.

**Fig. 3:**
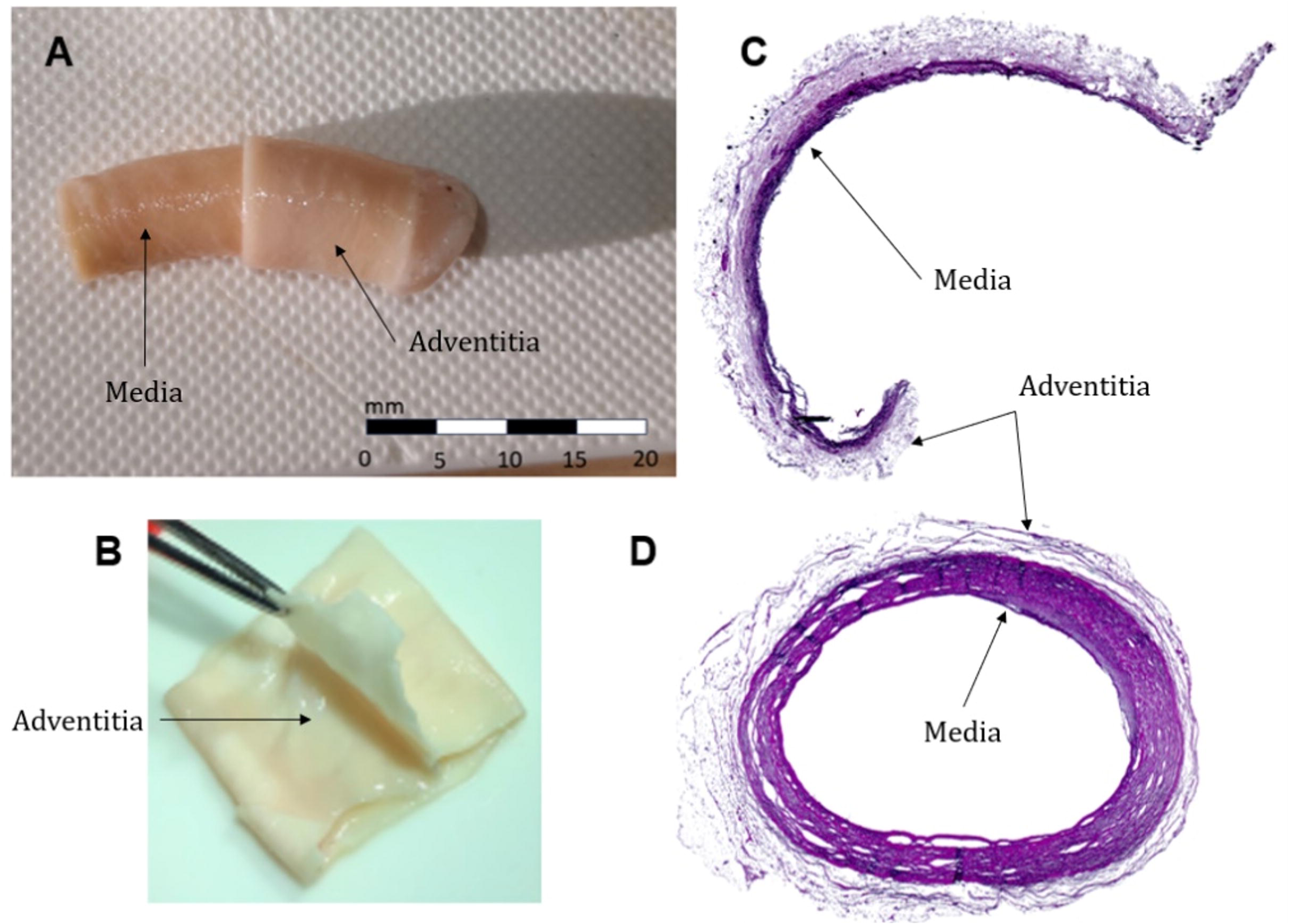
Opening angle [°] for both MI and A layers as well as intact ring I (in experimental protocol 1).

### Curvature

Curvature of the inner boundary is summarized in Fig. 4. The trends are opposite to those of the opening angles because of their inverse relation. Similarly to the opening angles, comparison of the final values (after 22 hours) revealed no statistically significant differences for both MI and A layers between the two experimental protocols, thus their values were merged. The curvature was 0.243±0.105 mm^−1^ for the MI layer, 0.343±0.071 mm^−1^ for the A layer, and 0.237±0.080 mm^−1^ for the intact wall (protocol 1 only). Similar results were observed for the outer boundary with curvature values of 0.237±0.145 mm^−1^, 0.280±0.058 mm^−1^ and 0.181±0.056 mm^−1^ for the MI and A layers and the intact artery wall, respectively.

**Fig. 4:**
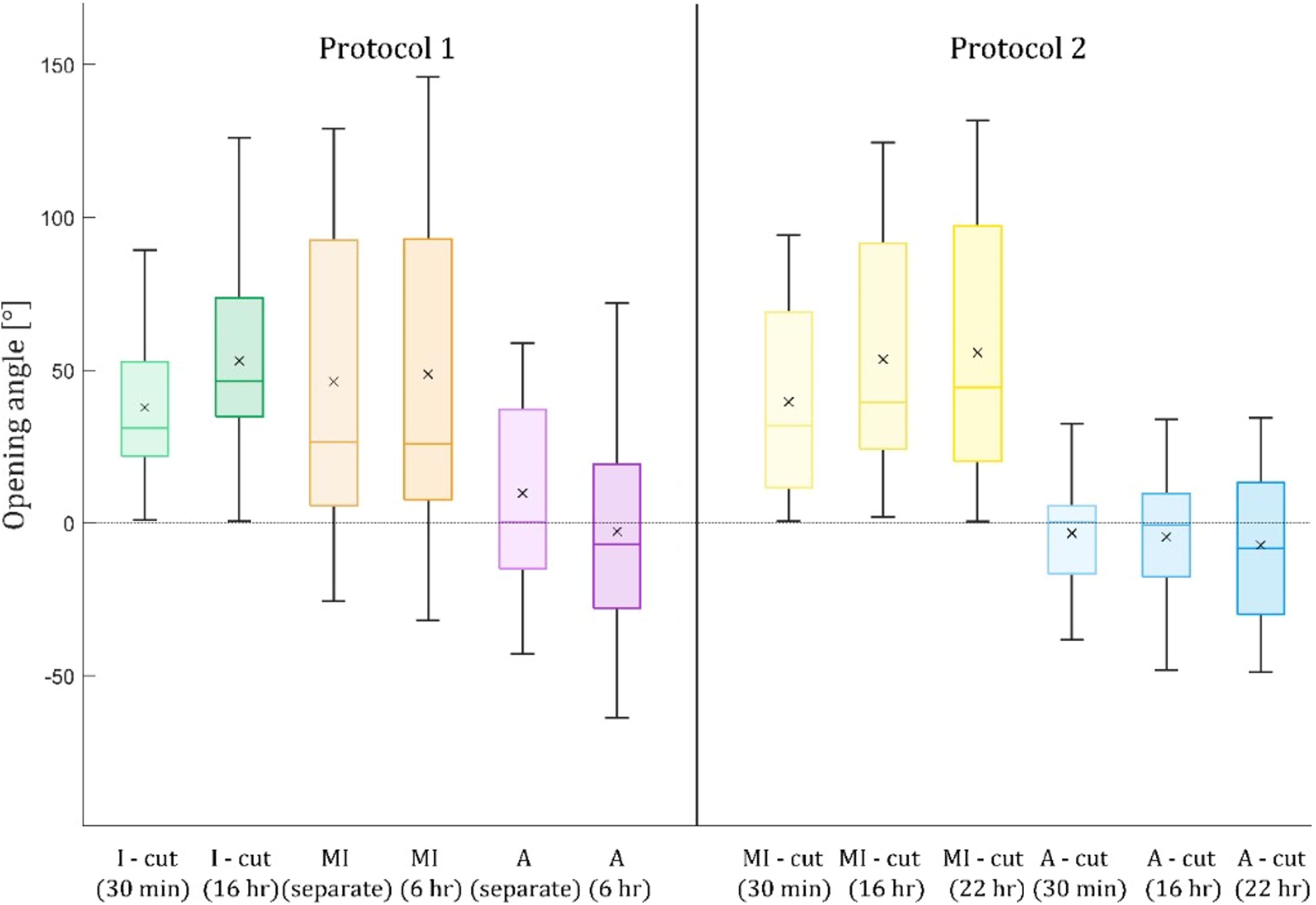
Curvature [mm^−1^] for both experimental protocols (protocol 1 is on grey background) – inner boundary (top), outer boundary (bottom).

### Impact of imperfect separation

Visual inspection, histology analysis and thickness measurement revealed varying amounts of media remaining on the adventitia layer (see Fig. 2C). The MI/A thickness ratio approximately quantifies this portion, with lower values indicating the adventitia layer more polluted with residual media. A positive correlation (ρ = 0.625; p < 0.005) was found between the adventitia curvature and the thickness ratio (see Fig. 5), suggesting that lower curvatures (i.e. positive opening angles) of adventitia are linked to a higher portion of residual media. Conversely, for a lower amount of residual media, opening angles were mostly negative according to their negative correlation with the thickness ratio (ρ = −0.432, p < 0.005). For the MI layer, trends were opposite to those found for adventitia, negative for the curvature (ρ = −0.419, p = 0.011) and positive for the opening angle. However, this last correlation was not statistically significant (ρ = 0.273, p = 0.069), although the p-value was close to the significance level of 0.05.

**Fig. 5:**
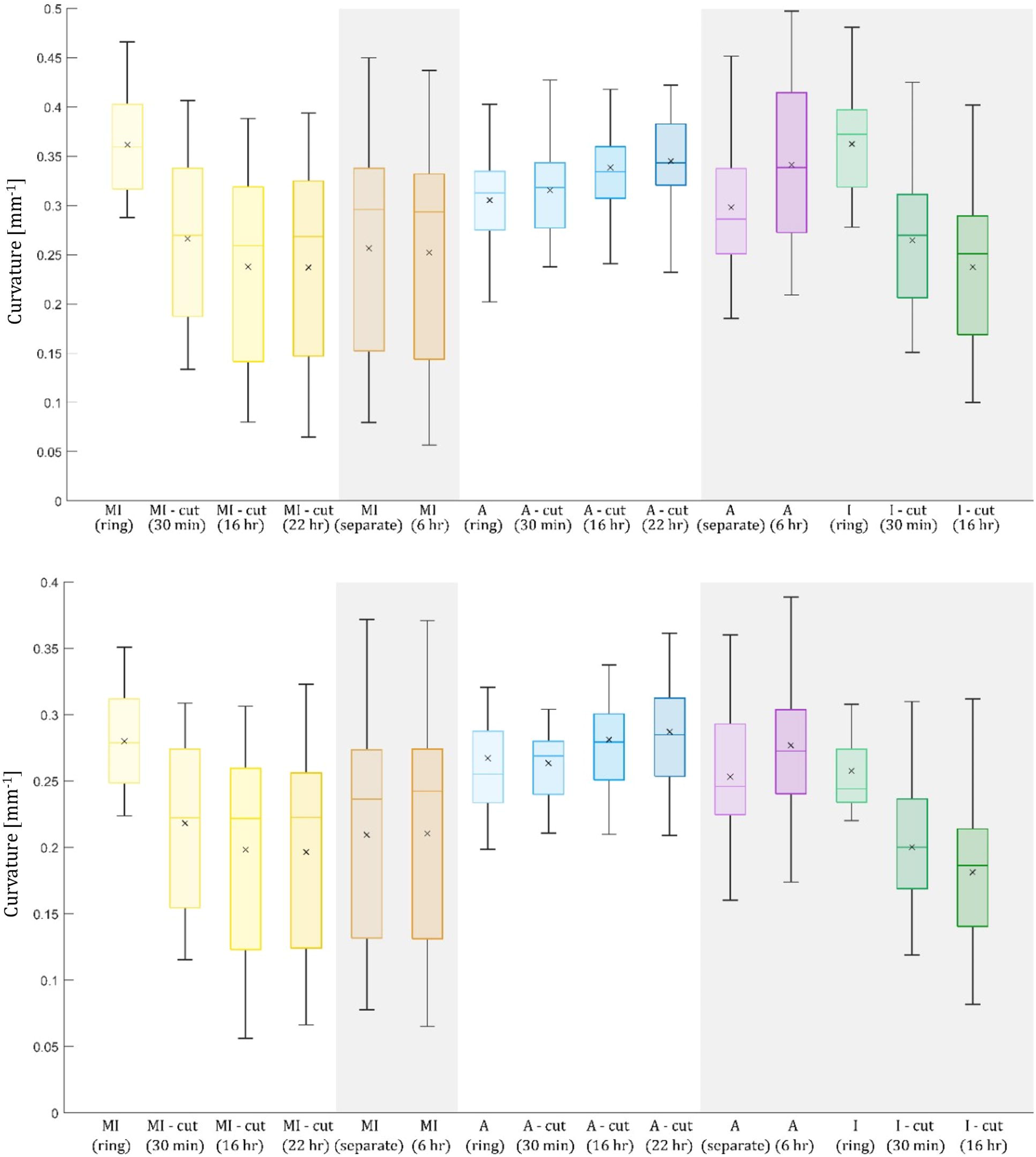
Correlation between thickness ratio and the opening angle (left) and inner curvature (right). The higher the thickness ratio, the lower portion of media remains on the adventitia layer.

### Length of specimens

Time development of the length of specimens in the circumferential direction at their inner and outer boundaries is presented in Fig. 6. Comparisons of inner and outer boundary lengths did not reveal significant differences between experimental protocols 1 and 2 thus, analogous to curvature and opening angle, the data was merged. The mean length of the inner boundary in the final stage was 18.404±2.703 mm for media, 19.654±2.280 mm for adventitia and 18.471±2.754 mm for the intact wall (protocol 1 only). For the outer boundary, the mean values were 21.845±2.387 mm, 24.205±2.983 mm and 23.869±2.695 mm for media, adventitia, and intact wall, respectively.

**Fig. 6:**
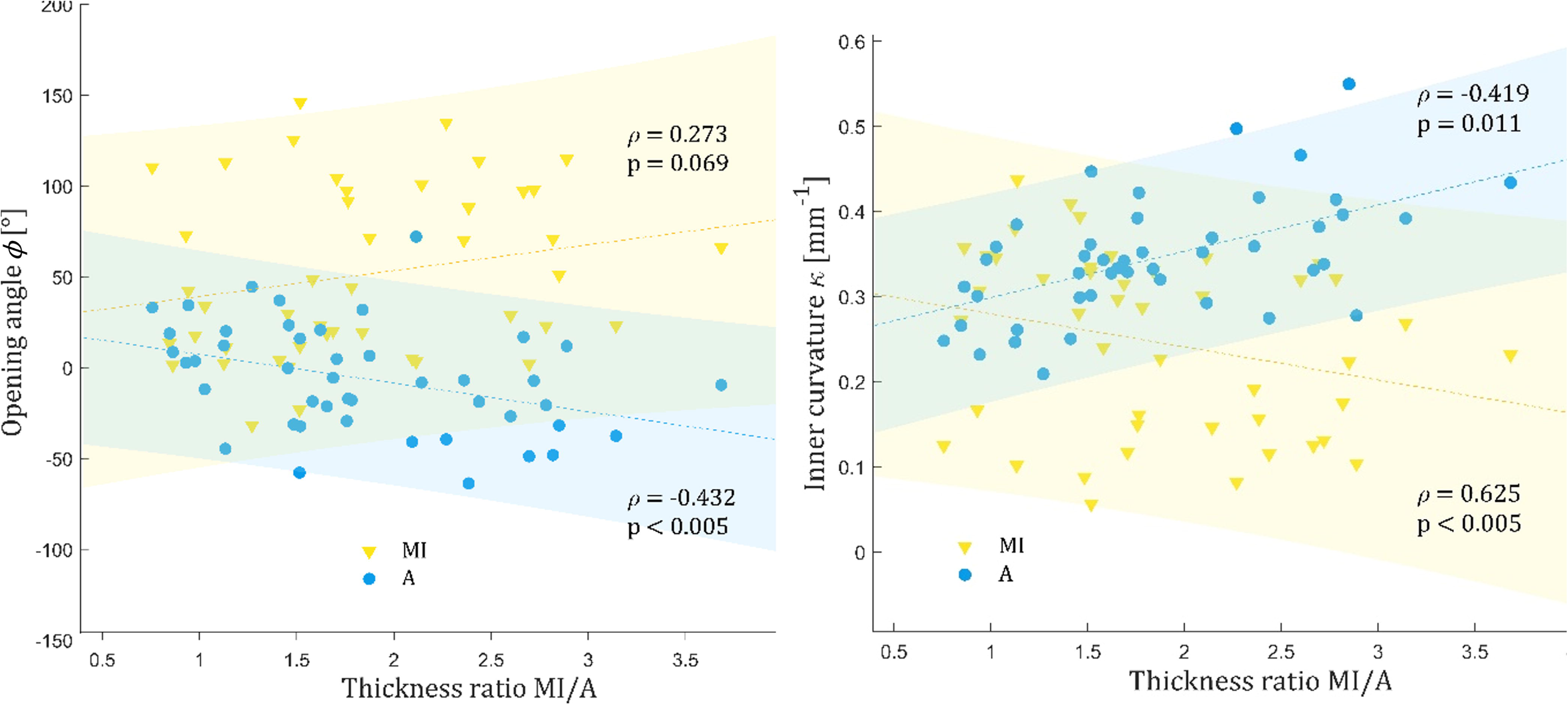
Length [mm] of the specimens for both experimental protocols (protocol 1 is on grey background) – inner boundary (top), outer boundary (bottom).

In addition, comparison between the length of the initial uncut ring and the length of the final state (after RS release) was conducted for each specimen using a paired t-test (for MI and A layers in protocol 2 and for I in protocol 1). The test revealed significant differences in lengths for all cases (MI, A, and I) at the inner boundary, while at the outer boundary, only A and I exhibited statistically significant differences. The inner boundary of the MI layer elongated by 6.12%, while the outer boundary remained unchanged. For the A layer, both the inner and outer boundaries shortened by 3.80% and 3.81%, respectively. The inner boundary of I elongated by 6.18%, whereas the outer boundary shortened by 4.18%. While the strain distribution in the media showed mostly a bending nature, in the adventitia, both bending and stretching were evident.

## 4. Discussion

Knowledge on the mechanical behaviour of arterial wall is essential for its computational modelling, where RSs are generally accepted to play a significant role, and their inclusion in computational models is often recommended. However, experimental evidence, especially in specific carotid artery layers and atheromatous arteries, remains sparsely explored and contradictory. Existing studies are based either on endarterectomy carotid samples (38), which exhibit small opening angles, or on coronary arteries (21), characterized by large opening angles. However, specimens from endarterectomy show much higher degree of atherosclerosis and incorporate atheroma instead of adventitia (and partially media), thus the results in (39) correspond rather to our MI layer. The same models and approaches were used for mouse samples in (40), recommending (together with (41)) the use of circumferential RS when analysing healthy and diseased arterial tissue. However, images of a closed segment published in (41) show some buckling effects causing unrealistic penetration of the fibrous cap (FC) or even lipid core into the lumen area. These effects were not discussed, although they indicate large negative stresses in the FC instead of positive stresses expected at FC rupture. This contradiction requires further experimental and computational investigation of this phenomenon.

In this study, two experimental protocols evaluating RS were compared. The first experimental protocol, echoing the methodology employed in (6), involves layer separation 16 hours after the radial cut, as opposed to the second experimental protocol introduced in this study, which opts for cutting the rings after the separation of layers. When the radial cut is made first and the separation of layers follows, the initial RS release happens while the layers are still joined together and influence each other. However, no statistically significant difference was found between final stages of both experimental protocols (after 22 hours) for any of the investigated parameters (thickness; opening angle; inner and outer curvature; inner and outer length) for both A and MI layer. Differences are in the attainable results: while the first method mediates insights into the intact wall, the second unfolds layer-specific deformation, which is of dominantly bending character in the intact wall as well as in the MI layer. However, the A shows much lower bending stiffness (it is proportional to the third power of the layer thickness) and much lower average opening angle values (or change of curvature)., Consequently, the adventitia deformation is dominated by contraction with a nearly constant average value of 3.8 % throughout the thickness and considering this effect on the RS distribution may be more important than the opening angle (or change of curvature) of the A layer.

When comparing adventitia and media, both layers show statistically significant differences for all the six parameters. The thickness was found to be 0.60±0.14 mm for the MI and 0.36±0.11 mm for the adventitia. These results corroborate that adventitia is much thinner than the MI layer. Nevertheless, our histology analysis revealed that the adventitia separation was not perfect (Fig. 2C) although the thicknesses are comparable with the values (0.7±0.13 mm and 0.47±0.07 mm, respectively) published in (24). As our MI/A thickness ratio was even higher than in that study, it suggests neither their separation was perfect.

The tendency of adventitia to negative angles (closing rather than opening, see Fig. 7 C, H) is not yet thoroughly described in literature. For instance, Teng et al. (39), although focused mainly on mechanical characterization (strength and stress-strain response) of both layers, mentioned the opening effect for the whole wall, as well as for the separated layers. However, their provided photo documentation is ambiguous, one figure shows an opened adventitia segment while another shows its zero opening. As they reported equal thickness of MI and A, their results correspond to ours with the lowest MI/A thickness ratio of approximately 1 (i.e. a significant portion of residual media on the adventitia), which gave also nearly zero angles in our study. It is also unclear whether their figures were taken in the solution or in the air, thus possibly influencing the results. Kural et al. (42) focused mainly on biaxial tensile tests but briefly mentioned RS characterized by the opening angle. A positive mean opening angle of 63° was reported for intact carotid artery walls while for separated layers some retraction or expansion was mentioned without being specified or further discussed. Esmaeili Monir et al. (18) focused on finite element modelling but showed one CCA sample with separated adventitia, media and intima. Here, a negative opening angle of −13° was recorded for adventitia which confirms our results.

**Fig. 7:**
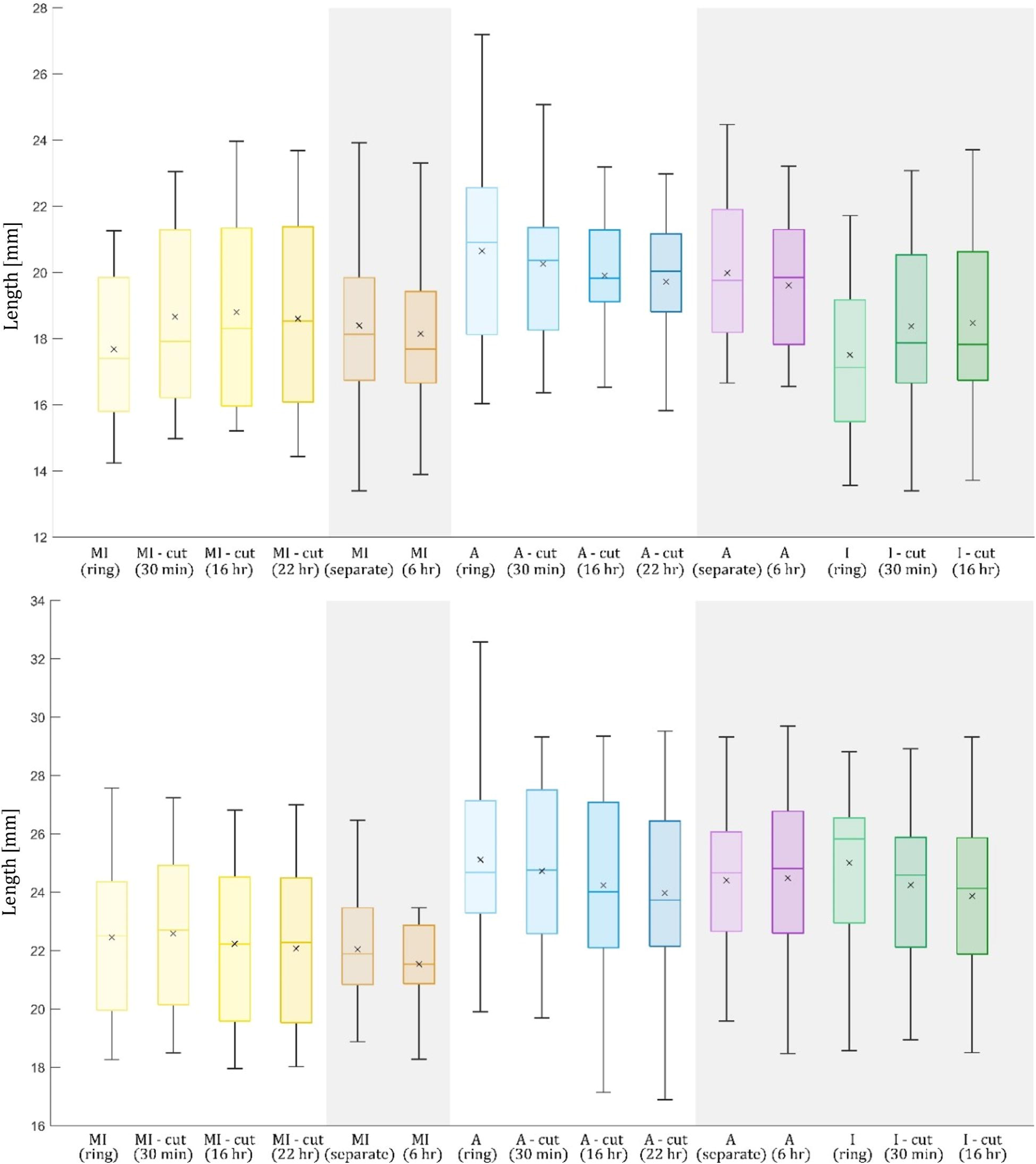
Example of typical opening of segments glued pointwise at the outer surface to a plastic cylinder. A, B: Intact wall – ring and after 16 hours; C, D: Adventitia and MI – 6 hours after separation, respectively; E, F: MI – ring and after 22 hours; G, H: Adventitia – ring and after 22 hours.

Sommer et al. (24) presented a decrease of curvature (opening tendency) in both MI and A layers of CCA immediately after their separation while some tendency to its re-increase with time occurred. They stated a slight increase of adventitia curvature (mean values) within 6 hours after separation, supporting thus partially the results of our study, in which we also accepted their approach applying the average curvature as RS quantifier. Although the final curvature was slightly higher than that of the intact wall, it was still much lower than the curvature of the uncut ring and represented thus an opening behaviour of adventitia. However, the specimens were glued on the inner surface to a plastic cylinder with diameter comparable to their inner diameter, thus a larger closing of the ring (negative opening angle) was precluded, and no statistical analysis was presented either. Statistical test of our results comparing the curvature of the inner adventitia boundary (in the stable state of experimental protocol 2) with the uncut adventitia ring confirmed the closing behaviour of adventitia indicating a significant release of RS. The same test done for the outer adventitia boundary has not reached statistical significance, because the remaining connective tissue makes the boundary blurry. Moreover, it induces some friction and constrains thus mutual movement of the free ends. This behaviour was noted when the free ends were mutually pushed together, distorting thus the circularity of the ring; the friction prevented the segment from further closing even though such tendency was apparent.

When comparing the length of the uncut adventitia ring and the length of the stable state cut specimen, the adventitia strips shortened significantly. This behaviour was also supported by the results from the intact wall specimens; here the statistical test revealed a difference between the inner and outer boundaries. The inner boundary (intima) elongated by 6.18%, whereas the outer boundary (adventitia) shortened by 4.18%. When comparing also the results from the separated MI layer, which elongates on the inner boundary but keeps the same length of the outer boundary, the following conclusions can be drawn: the intima on the inner surface tends to elongate and, conversely, the adventitia layer shows shortening behaviour. The experimental evidence on the length changes is scarce in literature; (18) reported elongation by 2.6% for media, 10.5% for adventitia and shortening by 9.3% for the intima layer for one CCA specimen. It is in contradiction to our results; however, it seems these values were read shortly after the radial cut; thus, they cannot represent the stable-state values reached much later. Additionally, these layer-specific length values were compared with the lengths of different intact specimen taken from the same CCA which may cause a deviation in the results. Sommer et al. (24) reported an axial shortening of the adventitia which was also observed in our experiments but could not be investigated and quantified due to small dimension of the specimens; in contrast, shortening in the circumferential direction is not discussed there. Holzapfel et al. (6) found that adventitia shortens significantly in both circumferential and axial directions after the release of RS, and the intima showed elongation, which corroborates our results although the media was found to shorten in the circumferential direction. Nevertheless, that study dealt with abdominal aortas differing significantly in composition and portion of the media layer from the muscular CCA, which may explain these differences.

The length modification was examined only as the uncut ring vs. the cut stable state (in protocol 1 for the intact wall and protocol 2 for the A and MI) allowing us a direct comparison of each sample and thus eliminating the interpatient variation. Surprisingly, when comparing the outer length of MI and the inner length of the A layer (for both separated rings before radial cut), the adventitia is always shorter (p-value for paired t-test <0.05) with values of 22.38±2.57 mm and 20.52±3.03 for MI and A, respectively. Although the layer separation of MI and A in the “turtleneck” fashion should not influence the bending RS, the tensile/compressive RS are influenced. This means the shortening effect of the A layer reaching nearly 10% strain may be even more significant.

The circumferential shortening of the adventitia documented in our study may also cause the curvatures of the intact specimen to be smaller than those of media, meaning the intact wall shows bigger opening angle than any of its components. The shortening tendency of the adventitia pulls the ends of the intact specimen further away from each other, resulting in a larger opening compared with the MI layer. This difference contradicts both hypotheses of constant strain and constant stress in the artery wall (throughout its thickness) and may change completely the RS distribution calculated in biomechanical modelling of artery wall. This impact should be verified in a future study.

## 5. Limitations

Acquisition of human tissue is quite challenging, so the sample number is limited. To increase the number of specimens, we were able to obtain multiple specimens from one sample. However, consideration of these specimens may induce a bias due to the natural inter-patient variability. Therefore, all the statistical analyses were recalculated on smaller data sets containing only one specimen per patient to avoid this bias, which allowed the use of paired t-test for direct specimen to specimen comparison (Wilcoxon signed-rank test as the nonparametric alternative). The results corroborate the conclusions obtained from the full datasets; no statistically significant difference was found between two different experimental protocols (p<0.05) for both MI and A layers when comparing all the parameters of interest: thickness, opening angle, inner and outer curvature, inner and outer length; the choice of experimental protocol does not have significant effect. Additionally, the length change after radial cut was confirmed on the reduced data as well. The comparison of length of inner boundary of A with outer boundary length of MI was confirmed on the reduced data set as well; the adventitia was 7.1 % shorter. The study is also limited by using circumferential specimens only, thus further measurements of axial residual stresses could enhance the description of the 3D residual stress distribution.

## 6. Conclusion

The experimental evaluation of the opening angles, curvatures and length changes of circumferential segments of human common carotid arteries and their layers revealed surprisingly increasing curvatures (negative opening angles) for separated adventitia layers. The inner MI layers showed an expectable decrease of curvature (i.e. positive opening angles), as well as the segments of intact (unseparated) artery wall, both dominated evidently by their bending. Together with significant shortening of adventitia after layer separation, these effects may completely change the distribution of stresses throughout the wall thickness. Neither of the hypotheses of constant strain or stress throughout the wall thickness, nor the residual stresses calculated based on opening angle of the artery wall segment are capable to describe the distribution of residual stresses correctly.

## Acknowledgements

This publication was supported by the Czech Science Foundation, research project No. 21-21935S, as well as by the project “Mechanical Engineering of Biological and Bio-inspired Systems”, funded as project No. CZ.02.01.01/00/22_008/0004634 by Programme Johannes Amos Commenius, call Excellent Research and Brno Ph.D. Talent Scholarship – Funded by the Brno City Municipality.

